# Model Based Inference of Large Scale Brain Networks with Approximate Bayesian Computation

**DOI:** 10.1101/785568

**Authors:** Timothy O. West, Luc Berthouze, Simon F. Farmer, Hayriye Cagnan, Vladimir Litvak

## Abstract

Brain networks and the neural dynamics that unfold upon them are of great interest across the many scales of systems neuroscience. The tools of inverse modelling provide a way of both constraining and selecting models of large scale brain networks from empirical data. Such models have the potential to yield broad theoretical insights in the understanding of the physiological processes behind the integration and segregation of activity in the brain. In order to make inverse modelling computationally tractable, simplifying model assumptions have often been adopted that appeal to steady-state approximations to neural dynamics and thus prevent the investigation of stochastic or intermittent dynamics such as gamma or beta burst activity. In this work we describe a framework that uses the Approximate Bayesian Computation (ABC) algorithm for the inversion of neural models that can flexibly represent any statistical feature of empirically recorded data and eschew the need to assume a locally linearized system. Further, we demonstrate how Bayesian model comparison can be applied to fitted models to enable the selection of competing hypotheses regarding the causes of neural data. This work establishes a validation of the procedures by testing for both the face validity (i.e. the ability to identify the original model that has generated the observed data) and predictive validity (i.e. the consistency of the parameter estimation across multiple realizations of the same data). From the validation and example applications presented here we conclude that the proposed framework provides a novel opportunity to researchers aiming to explain how complex brain dynamics emerge from neural circuits.

## 1 Introduction

Networks of interconnected neural ensembles form the substrate upon which highly structured neural activity unfolds in a way which is thought to facilitate the integration and segregation of information in the brain (Sporns, 2013; Varela et al., 2001). Large scale neuronal models (Breakspear, 2017; Deco et al., 2015) provide a way of understanding mechanistically how these dynamics arise from the interaction of: (a) the *functioning* of neurons as is dictated by their intrinsic biophysical properties; and (b) the *structure* of the network that connects them. Typically, functional integration across brain networks has been measured in terms of functional connectivity (i.e. assessing statistical dependencies between distinct neuronal activities). However, these purely statistical approaches lack any mechanistic description as to how integration may arise. Better characterisation of mechanisms in terms of known structure and function of neuronal circuits can give insight as to how segregation and integration are achieved in the brain (Deco et al., 2015). The construction of mathematical models embodying the essential dynamics and structure to describe a particular phenomenon provides a theoretical framework by which to derive new testable hypotheses of the mechanisms that generate large scale brain activity such as that measured in the electroencephalogram or local field potentials. The tools required to identify a model (and its parameters) from a set of experimental data stem from developments in the *inverse* modelling of biological systems. These tools allow for an experimenter to work backwards from a set of noisy and often incomplete empirical observations to infer the causal factors that produced them.

Whilst powerful, the optimization algorithms used in the computation of inverse models typically involve a parameter search that requires numerous evaluations of a given model. To render inference computationally efficient, methods often take steps to reduce the complexity of the models to be inverted. For instance, Dynamic Causal Modelling (DCM), a common method for inverse modelling of neuroimaging data, fits directly to the cross- and auto-spectra of a time series by invoking simplifying assumptions as to the model’s dynamics i.e. that the measured activity is generated from a system close to equilibrium with dynamics that are approximately linear. By making this assumption it is possible to approximate the system’s behaviour with a single transfer function that can be used to predict the spectral response of a system to innovations with a specific form (Friston et al., 2012). This simplification of candidate models’ dynamics effectively removes a large subset of dynamical regimes in which the neural system could potentially operate, making the problem easier to solve (as equations no longer need to be numerically integrated), but effectively restricts the types of behaviours that may be explored.

In this paper we compile a framework for the inference of brain networks from neural data that aims to avoid the requirement to simplify model dynamics. This approach follows on from DCM but uses an optimization scheme based upon Approximate Bayesian Computation (ABC) that makes it possible to invert neuronal models without invoking assumptions of local linearity. ABC is a Bayesian optimization procedure dependent on sampling (Beaumont et al., 2002). It is a “likelihood free” algorithm (Marin et al., 2012) making it well suited for complex models for which there is a large state and/or parameter space, exhibit stochastic or nonlinear dynamics, or require numerically expensive integration schemes to solve. The method has been successfully employed and validated across several domains of systems biology (Excoffier, 2009; Liepe et al., 2014; Toni and Stumpf, 2009; Turner and Sederberg, 2012), but has not yet seen wide usage in large scale neuroscience. Usefully, the schema allows models to be flexibly inverted upon data features beyond that of the auto- and cross-spectra; in the examples shown here we use Non-Parametric Directionality Analysis (NPD)-an estimator of directed functional connectivity between time series. Further, because of the relaxation of assumption on model dynamics, this scheme opens up inverse modelling to exploring neural circuits that can describe more complicated behaviours such as itinerancy/metastability (Deco et al., 2017) or the role of factors such as transmission delays and noise (Deco et al., 2009) by explicitly incorporating them into a model.

Specifically we perform parameter estimation using an algorithm based on sequential Monte Carlo ABC (ABC-SMC; Toni et al., 2009) that is well suited for use with the types of data and models typically used in studies of large scale brain activity. We use an adaptation of this scheme that incorporates kernel estimation of parameter marginals and their dependencies using copula theory (Li et al., 2017) to make it better suited to highly parameterized large scale neuronal models. We cast all examples presented here in the context of beta band (14-30 Hz) dynamics found in the Parkinsonian cortical-basal ganglia-thalamic circuit using a previously reported model (van Wijk et al., 2018) and experimental data (West et al., 2018). In the present work we will retain steady state data features (spectra and directed functional connectivity) as the basis on which to invert models of connectivity but use a framework that does not restrict model dynamics. Thus, complex features such as intermittency in oscillations (e.g. Parkinsonian beta bursts) contribute to the model output and are open to exploration in post-hoc simulations of the fitted system (c.f. West et al., 2020). We first examine the properties of the inversion scheme, investigating parameter estimates and convergence. Further, to validate this schema we use a similar approach to that used previously in the validation of methods such as DCM by first testing the so called *face* validity to examine whether the method is able to recover and reidentify parameters of the model that has caused the data (Moran et al., 2009). We subsequently provide a test of *predictive* validity by examining whether the method can provide consistent estimation of parameters across multiple realizations of the data. Finally, we demonstrate how the scheme can be scaled up to be applied to a large model space with potential to test a set of biologically relevant hypotheses.

## 2 Methods

### 2.1 Overview of Sequential Monte Carlo Approximate Bayesian Computation for Inverse Modelling of Neural Data

We present an overview of the framework using ABC-SMC and its adaptations for applications to large scale neural models is figure 1. We summarise the previous methodologies that we synthesise in this work in supplementary table I. The algorithm takes a form in which several processes are repeated multiple times within their parent process (figure 1; inset). The schema is contingent on simulation of pseudo-data by a generative forward model – a description of the neural dynamics - (figure 1A; green box) given a set of proposal parameters sampled from a prior (Gaussian) distribution (figure 1C; turquoise box). This pseudo-data can then be compared against the empirical data by first using a common data transform (i.e. a summary statistic of the data) and then assessing their similarity by computing the objective function (goodness-of-fit R^2^) (figure 1B; blue box). This model fit provides a measure by which parameter samples are either rejected or carried forward depending on a threshold on the goodness-of-fit, in order to generate the next proposal distribution in the sequence. When the process in figure 1C is iterated with a shrinking tolerance schedule, ABC can be used to approximate the posterior parameter distribution at convergence (figure 1C; orange box). Finally, if the process described above is repeated over several competing models then the approximate posterior distribution may be used to assess each model’s fitness via model comparison (figure 1D; purple box). The exact details of each process outlined in the figure are given below.

**Figure 1.**
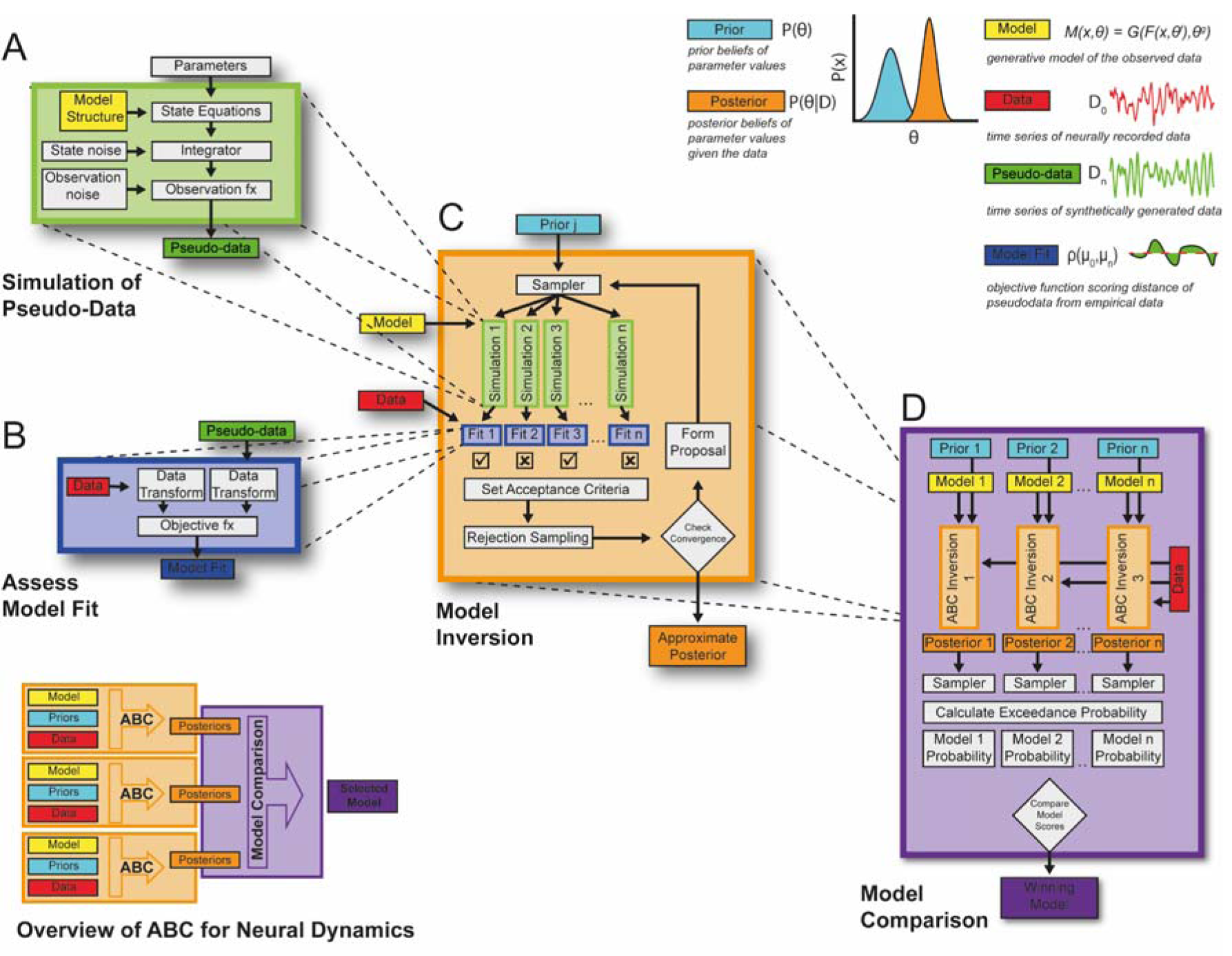
Framework for application of Approximate Bayesian Computation for model based inference of brain network dynamics. **(Inset)** This schematic gives an overview of the structure of the framework described in this paper. Individual generative models are specified as a set of state equations (yellow boxes) and prior distribution of parameters (light blue boxes) that will be used to approximate the posterior density of the parameters (orange boxes) of the system generating the observed data (red boxes) with varying degrees of fit (blue boxes) using ABC. The approximate posterior distribution can then be used to compare models and decide on a winning model or family of models (purple box). **(A) Generation of pseudo-data** by integrating state equations parameterized by a sample of parameters drawn from a prior or proposal distribution. Models can incorporate stochastic innovations as well as a separate observation model to produce samples of pseudo-data (green boxes). **(B) Pseudo-data is compared against the real data** using a data transform common to both that provide a summary statistic of the time series data (i.e. spectra and functional connectivity). The simulated and empirical data are then compared by computing the objective function that can be used to score the model fit (blue boxes). **(C) ABC sequentially repeats the processes in boxes A and B** by iteratively updating a proposal distribution formed from accepted samples. Samples are rejected depending on an adaptive thresholding of the objective scores with the aim to reduce the distance between summary statistics of the data and pseudo-data. This process iterates until the convergence criterion is met and the proposal distribution is taken as an approximation of the posterior distribution. **(D) By repeating the ABC process in box (C) over multiple models**, the approximate posteriors can be used to evaluate the model probabilities. This process samples from the posterior many times to compute the probability of each model exceeding the median accuracy of all models tested. This “exceedance” probability can then be used to compare the model’s ability to accurately fit the data and select the best candidate model given the data.

### 2.2 Generative Model of Neural Pseudo-Data

The fitting algorithm is based upon sampling from a sequence of proposal distributions over parameters to generate realizations of the generative process that we refer to as pseudo-data (figure 1A; green box). A model *M* is specified by the state equations of the dynamical system *F* and observation model *G*:

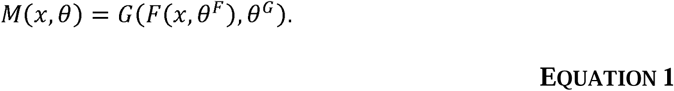

The state equations of the model *F* describe the evolution of states *x* with parameters *θ*^*F*^. This model describes the underlying dynamics that give rise to the evolution of the neural states. The observation model *G* describes the effects upon the signals that are introduced by the process(es) that occur during experimental acquisition and recording of the neural signal(s) *x* and is parameterized by *θ*^*G*^. This observation model aims to account for confounds introduced beyond that of the generative model of the data such as changes in signal-to-noise ratio, crosstalk, or other distortions of the underlying generative process.

In general, model *M* could describe any dynamical system describing the time evolution of neural data such as spiking networks, conductance models, or phenomenological models (e.g. phase oscillators, Markov chains). In this paper we use a model based upon the anatomy of the cortico-basal-ganglia-thalamic network that is constructed from coupled neural mass equations (David and Friston, 2003; Jansen and Rit, 1995), the biological and theoretical basis of which has been described previously (Marreiros et al., 2013; Moran et al., 2011; van Wijk et al., 2018). We adapt the original equations to explicitly incorporate stochastic inputs and finite transmission delays. This system of stochastic delay differential equations is solved using the Euler-Maruyama method. For details of the modelling, as well as details of the integration of the state equations please see Supplementary Information II. The full set of state equations are given in supplementary information III. For results sections 3.1 through to 3.3 we use a subnetwork of the full model comprising the reciprocally coupled STN/GPe. This model can be again divided into separate models (for the purposes of performing face validation and example model comparison) by constraining priors on the connectivity between the STN and GPe. Section 3.4 uses a wider model space comprising a set of systematic variations upon the full model (see figure 5).

Parameters of both the generative and observational model can either be fixed or variable. In the case of variable parameters (parameters to be fit), a prior density encoding *a priori* beliefs about the values that the parameters take must be specified. This is encoded through a mean and variance for each parameter, with the variance encoding the inverse-precision of a prior belief. In this way fixed parameters can be thought as being known with complete confidence. We take the majority of prior values for parameters of the cortico-basal-ganglia-thalamic network from (van Wijk et al., 2018) but set some delays and connection strengths given updated knowledge in the literature. A table of parameters can be found in Supplementary Information IV.

### 2.3 Model Inversion with Sequential Monte Carlo ABC

#### 2.3.1 Algorithm Overview

In order to estimate the parameters of the model *M* given the information provided by the empirical recordings we use an algorithm based upon ABC-SMC (Del Moral et al., 2012; Toni et al., 2009). Most generally, the algorithm forms a sample of draws (particles) taken from a prior distribution and then goes on to estimate an intermediate sequence of proposal distributions via iterative rejection of the particles. Given a suitable shrinking tolerance schedule, the simulated pseudo-data (generated from the sample-parameterized forward model) and the empirical data should converge as the proposal distribution approaches the true posterior.

The ABC algorithm is illustrated in figure 1C (orange box) and follows the procedure below. Probability densities are given by *P*(·); parameters are indicated by *θ*; models by *M*; data by *D;* and distances by ρ. Samples are indicated by hat notation (i.e. ^); subscripts indicate the sample number; proposal distributions are indicated by an asterisk (i.e. *P*(·)*); and subscripts equal to zero denote belonging to the empirical data (i.e. *µ*_1_ and *µ*_0_are summary statistics of sample 1 and of the empirical data respectively).

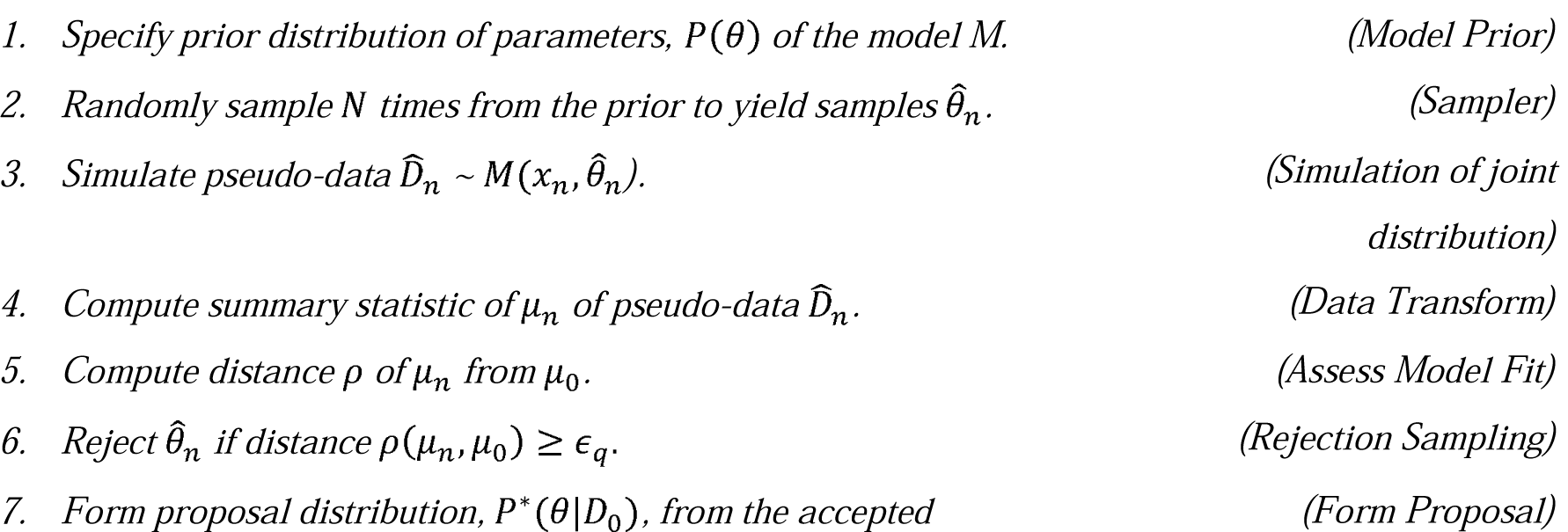
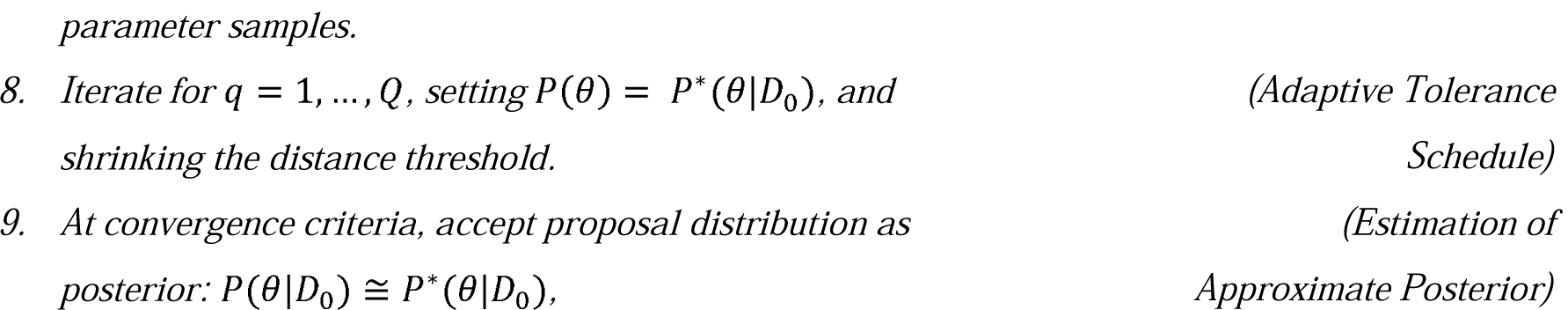

To avoid sample wastage across iterations, we store the samples from step (2) and their resulting distance from the data (step 5) across the *Q* iterations such that they are propagated through a sequence of intermediate distributions. In this way, at step (7) the updated proposal distribution comprises samples from both current and past draws selected on the basis of a threshold calculated over the current draw.

We estimate the “distance” of the data from pseudo-data using the coefficient of determination (R^2^) as an objective function ρ(*µ*_*n*_,*µ*_0_) and closely related to the mean squared error (MSE). We note that in nonlinear models (and linear models without an intercept), R^2^ takes value in [-∞ 1] with 1 indicating a perfect fit, and negative R^2^ denoting a fit worse than that of the mean of the data (Cameron and Windmeijer, 1997). Setting a shrinking distance threshold ensures that the posterior estimates converge upon solutions that most accurately reproduce the summary statistics of the observed data (Del Moral et al., 2012). With non-negligible ϵ_Q_, the algorithm samples from an approximate posterior distribution 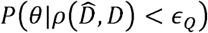 rather than the true posterior *P*(*θ*|*D*). Thus the upper bound on the error of parameter estimates is therefore determined by how far ϵ_Q_ is from zero (Dean et al., 2014).

#### 2.3.2 Adaptive Tolerance Schedule

To facilitate incremental sampling from a sequence of increasingly constrained target distributions we set an adaptive tolerance schedule. This is specified by determining a predicted gradient for the average distance of the next set of samples:

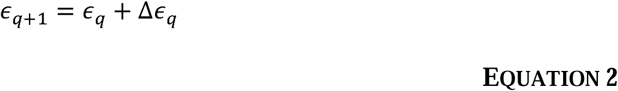

where the expected change in the distance of the new samples Δϵ_*q*_ is given by:

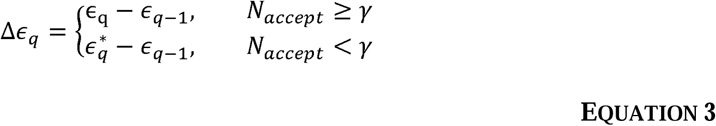

where *N*_*accept*_ is the number of accepted samples and γ is a minimum criterion on the accepted sample size to carry forward the tolerance shrinkage at its current gradient. If *N*_*accept*_ < γ then this gradient is assumed to be too steep and the expected gradient is recalculated using a modified tolerance 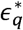 that is computed using the median distance ρ between the sample pseudo-data from that real (i.e.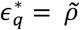, where ∼ indicates the median). Thus γ parameterizes the coarseness of the optimization. If γ is very large (e.g. >99% of *N*) then the algorithm will converge slowly but accurately, whereas if γ is very small (e.g. 1% of *N*) the algorithm will be inaccurate and biased. We set γ to be the two times the estimated rank of the parameter covariance matrix i.e.*rank*(Σ) (for details of estimation see bottom of section 3.3.3).

#### 2.3.3 Formation of Proposal Distributions

Following rejection sampling, the proposal density *P*^*^(*θ*|*D*_0_) must be sampled from the accepted parameters sets. We use a density approximation and copula approach for ABC similar to that described in Li et al., (2017). In the initial draw of samples from the prior we treat the data as Gaussian and form a multivariate normal:

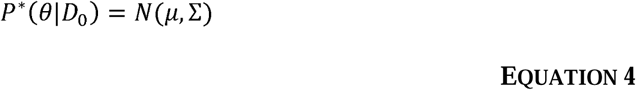

where *µ* is a vector of the univariate expectations and Σ their covariances. In subsequent iterations whereby a minimum sample is accumulated, we use nonparametric estimation of the marginal densities over each parameter using a non-parametric kernel density estimator (Silverman, 2003). This approach does not assume normality of parameter distributions and may estimate multimodal parameter distributions. This flexibility allows for sampling across multiple possible maxima at once, particularly at intermediate stages of the optimization. The bandwidth (determining the smoothness) of the kernel density estimator is optimized using a log-likelihood, cross-validation approach (Bowman, 1984).

We then form the multivariate proposal distribution using the t-copula (Nelsen, 1999). Copula theory provides a mathematically convenient way of creating the joint probability distribution whilst preserving the original marginal distributions. Data are transformed to the copula scale (unit-square) using the kernel density estimator of the cumulative distribution function of each parameter and then transformed to the joint space with the t-copula.

The copula estimation of the correlation structure of the parameter distributions acts to effectively reduce the dimensionality of the problem by binding correlated parameters into modes. The effective rank of the posterior parameter space (used in the computation of the adaptive tolerance schedule and reported in the results as a post-hoc assessment of parameter learning) can be estimated by taking the eigenvalues of the covariance matrix and normalizing the coefficients by their sum. Using a cumulative sum of the ordered coefficients we can then determine the number of modes that can explain 95% of the variance of the parameter samples.

#### 2.3.4 Priors on Model Dynamics

We aim to identify models yielding non-stationary, stochastic outputs and so we explicitly incorporate this into the prior specification of model dynamics. Concretely, we reject simulations in which there is either strong periodicity in the envelope of the signal, or if the envelope is divergent, respective signs of the analytic signal computed from the Hilbert transformed broadband signal *S*(*t*) = |(*x*(*t*) |. We of a signal that is either close to equilibrium or unstable. We define the envelope *S* as the magnitude compute envelope periodicity using the autocorrelation (scaled to unity at zero-lag) of this signal and remove signals with statistically significant correlations at lags ±2ms. To determine divergence, we test for monotonicity of the envelope by applying a non-parametric rank correlation test. In the case of a divergent signal, ranks will be significantly increasing or decreasing and so we remove signals with statistically significant correlation. Models failing these criteria are automatically removed by setting their R^2^ to -∞.

### 2.4 Model Comparison

In the process of model-based inference, hypotheses must be compared in their ability to explain the observed data. In this section we describe a method (Grelaud et al., 2009; Toni and Stumpf, 2009) in which models fit with ABC can be formally compared. At convergence ABC approximates the posterior parameter distribution *P*(*θ*|*D*_0_) that may be used to approximate the model evidence *P(M*|*D*_0_). This estimate is made by drawing *N* times from the posterior and then computing an exceedance probability to estimate the marginal probability of the *j*^*th*^ model *M* given data *D*_0_(the approximate model evidence):

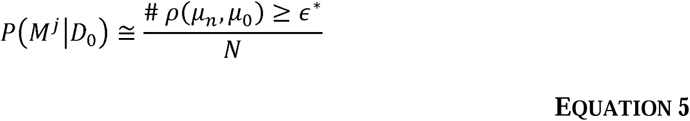

where ϵ* is a threshold on the distance metric ρ that is suitably small to give an acceptable fit on the data. If ϵ*is held constant and the data is identical between models, then the exceedance probabilities may be compared to yield the model that gives the most accurate fit. In practice we set ϵ* to be the median distance of all sets of models. In order to compare models a Bayes factor *K* can be constructed:

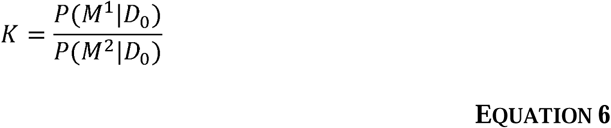

where *K* gives the strength of evidence of model 1 over model 2 that can be interpreted as proposed by (Kass and Raftery, 1995). In addition to calculating a Bayes factor derived from the estimated model evidences, we also provide a post-hoc examination of model complexity in the form of a divergence of the posterior from the prior (Friston et al., 2007; Penny, 2012). Specifically, we estimate the Kullback-Lieber divergence D_KL_ of the posterior density *P*(*θ*|*D*_0_) from the prior density *P(*θ*)* over *F* discretized bins of the density:

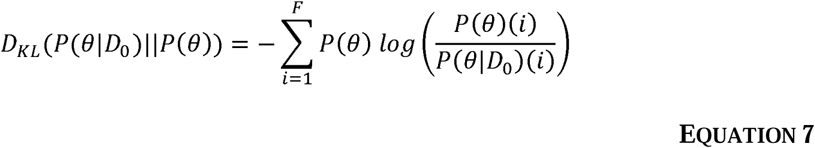

This is a simplification of the full multivariate divergence and ignores the dependencies between variables encoded in the posterior covariance. We use the full multivariate divergence (given in supplementary information V) that makes a Gaussian approximation to the marginals, whilst accounting for their covariance structure. This is a reasonable approximation to make as we only expect densities to be non-Gaussian in the proposal densities. Given that the algorithm has reached convergence, proposal densities over parameters will be normally distributed. The D_KL_ can then be used as a post-hoc regularization of the model comparison in which model accuracy provides the primary objective function, whilst D_KL_ provides a secondary discriminator to determine the degree of overfitting. When two models have identical accuracy, they can be further segregated by examining D_KL_ and favouring the model that deviates the least from the prior parameters as being the most parsimonious of the two explanations of the data since it requires less deviation from the experimenter’s prior expectations of parameter values (as an appeal to Occam’s razor; MacKay, 2003). Finally, we combine both the model accuracy and D_KL_ to yield a combined metric (accuracy-complexity score; ACS) accounting for both accuracy and complexity for model *j*:

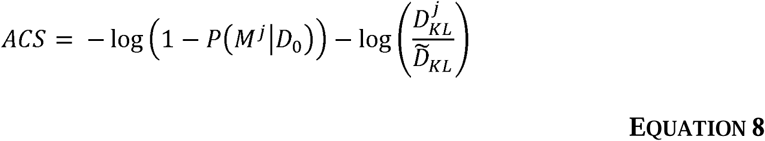

where 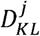 is the divergence of posteriors from priors for the *j*^*th*^ model 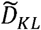 and is the median divergence across the whole model space. With increasing accurate and/or minimally divergent models the ACS metric will tend towards large positive values.

### 2.5 Empirical Data: Recordings from Parkinsonian Rats

Summary statistics are computed from empirical data and then used to fit the generative forward model. In the example implementation used in this paper we use multisite basal ganglia and single site cerebral cortex recordings in rats (n=9) that have undergone a 6-hydroxydopamine (6-OHDA) induced dopamine depletion of the midbrain, a lesion model of the degeneration associated with Parkinsonism in humans (Magill et al., 2006, 2004). The animals were implanted with two electrodes to measure local field potentials (LFP) from multiple structures in the basal ganglia: dorsal striatum (STR), external segment of the globus pallidus (GPe), and the subthalamic nucleus (STN). Additionally electrocorticography was measured over area M2 of the motor cortex, a homologue of the Supplementary Motor Area (SMA) in humans (Paxinos and Watson, 2007). Animals were recorded under isoflurane anaesthesia and during periods of “cortical-activation” induced by a hind-paw pinch (Steriade, 2000). The details of the experimental procedures were previously published (Magill et al., 2006, 2004). Experimental procedures were performed on adult male Sprague Dawley rats (Charles River) and were conducted in accordance with the Animals (Scientific Procedures) Act, 1986 (UK), and with Society for Neuroscience Policies on the Use of Animals in Neuroscience Research. Anesthesia was induced with 4% v/v isoflurane (Isoflo; Schering-Plough) in O2 and maintained with urethane (1.3g/kg, i.p.; ethyl carbamate, Sigma), and supplemental doses of ketamine (30 mg/kg, i.p.; Ketaset; Willows Francis) and xylazine (3 mg/kg, i.p.; Rompun, Bayer).

Pre-processing of time series data (LFP and ECoG) was done as follows: data were 1) truncated to remove 1 second (avoid filter artefacts); 2) mean corrected; 3) band-passed filtered 4-100 Hz with a finite impulse response, two-pass (zero-lag) with optimal filter order; 4) data were split into 1 second epochs with each epoch subjected to a Z-score threshold criterion such that epochs with high amplitude artefacts were removed.

### 2.6 Computation of Summary Statistics

We derive a set of summary statistics from signal analyses of the experimental and simulated time series. These statistics transform both the data and pseudo-data into the same feature space such that they can be directly compared (figure 1B; blue box). It is important to note that the summary statistic is vital in determining the outcome of the inverse modelling with ABC (Beaumont et al., 2002). The set of statistics must effectively encode all phenomena of the original data that the experimenter wishes to be modelled.

#### 2.6.1 Frequency Spectra

We use the autospectra to constrain the oscillatory characteristics of each neural mass population. Auto-spectral analyses were made using the averaged periodogram method across 1 second epochs and using a Hanning taper to reduce the effects of spectral leakage. Frequencies between 49-51 Hz were removed so that there was no contribution from 50 Hz line noise. 1/f background noise was removed by first performing a linear regression on the log-log spectra (at 4–48 Hz) and then subtracting the linear component from the spectra. This ensured that the inversion scheme was focused upon fitting the spectral peaks in the data and not the profile of 1/f background noise. To simplify observation modelling of differences in experimental recording gains between sites, all spectra were normalized by dividing through by their summed power at 4-48 Hz.

#### 2.6.2 Directed Functional Connectivity

To quantify interactions between populations, we use non-parametric directionality (NPD; Halliday 2015), a directed functional connectivity metric which describes frequency resolved, time lagged correlations between time series. The NPD was chosen as it directly removes the zero-lag component of coherence and so statistics are not corrupted by signal mixing. This has the effect of simplifying observation modelling by reducing the confounding effects signal mixing.

Estimates of NPD were obtained using the Neurospec toolbox (http://www.neurospec.org/). This analysis combines Minimum Mean Square Error (MMSE) pre-whitening with forward and reverse Fourier transforms to decompose coherence estimates at each frequency into three components: forward, reverse and zero lag. These components are defined according to the corresponding time lags in the cross-correlation function derived from the MMSE pre-whitened cross-spectrum. This approach allows the decomposition of the signal into distinct forward and reverse components of coherence separate from the zero-lag (or instantaneous) component of coherence which can reflect volume conduction. The method uses temporal precedence to determine directionality. For a detailed formulation of the method see Halliday, (2015); and for its validation see West et al., (2020b). We ignored the instantaneous component of the NPD and use only the forward and reverse components for all further analyses.

#### 2.6.3 Data Pooling and Smoothing

In all optimizations using empirical data in this study we used the group-averaged statistics computed from recordings from a group of unilaterally 6-OHDA lesioned animals. As a final processing step, both the autospectra and NPD were smoothed to remove noise as well as to reduce them to their main components (peaks). This was achieved via fitting of a sum of one, two, or three Gaussians to the spectra using the adjusted R^2^ as a guide for selecting the most appropriate sum of Gaussians. This allows the spectra to be reduced to a smooth sum of normal modes corresponding to the main peaks in the spectra (i.e. alpha (8-12 Hz), low beta (12-21 Hz) and high beta/gamma frequencies (21-30 Hz)). Empirical and simulated data were transformed similarly to produce equivalent autospectra and NPD. The simulated data did not undergo regression of 1/f background in the autospectra; nor Gaussian smoothing of the data features as the simulations aimed to approximate the smoothed data features as directly as possible. In both cases these additional transforms may be misappropriated by the optimization procedures to yield superficially close fits.

### 2.7 Validation of ABC Procedures for Parameter Inference and Model Identification

#### 2.7.1 Testing the Predictive Validity of the ABC Parameter Estimation with Multi-start

To test whether the ABC estimator will: a) yield parameter estimates that are unique to the data from which they have been optimized; and b) yield consistent estimation of parameters across multiple instances of the schema, we performed a multi-start. We performed two procedures of ten multi-starts for a single STN/GPe model (using identical priors). The two multi-starts were identical except for the data to which they were fitted. Over the evolution of each optimization, the posterior parameters are reported as the maximum a posteriori (MAP) estimates taken as the median of the marginal distributions.

When testing the point (a)-that parameter estimates are unique to the data from which they are fitted - we performed a 10-fold cross-validation procedure in which we used a one-sample Hotelling procedure to test for significant difference of each fold’s mean from that of the left-out sample. We report the probability of the folds that yielded a significant test, with high probability indicating that the left-out MAP estimates are likely to deviate from the rest of the fold. In this way we can identify the probability of an ABC initialization yielding a non-consistent sample. Secondly, we test (b) – that MAP estimates are unique to the data on which they have been fitted-using the Szekely and Rizzo energy test (Aslan and Zech, 2005) between the samples from data A and B, with the null-hypothesis that the multi-start samples derived from different data arise from the same distribution. Finally, we use a Multivariate Analysis of Variance (MANOVA) procedure to test for difference in means between the two multivariate samples. In the case of a positive MANOVA test, post-hoc analyses of differences in means of each parameter are made using two sample ANOVAs.

#### 2.7.2 Testing the Face Validity of the Model Comparison Procedures – Model Identification

To test the face validity of the model comparison framework, we constructed a confusion matrix, an approach commonly used in machine learning to examine classification accuracy. Three different models of the STN/GPe network were fit to the empirical data and then using the fitted parameters three synthetic data sets were simulated. We chose a model with reciprocal connectivity: (1) STN ↔ GPe; and then two models in which one connection predominated: (2) STN → GPe and (3) GPe → STN. The three models (with the original priors) were then fitted back onto the synthetic data. A model comparison procedure was then performed to see whether it could correctly identify the best model as the one that had generated the data. The model comparison outcomes (accuracy of the fit; D_KL_ of posteriors from priors; and the combined ACS measure) were then plotted in a 3 × 3 matrix of data versus models. In the case of valid model selection, the best fitting models should lay on the diagonal of the confusion matrix.

#### 2.7.3 Testing the Scalability of the Framework with Application to the Full Model Space

In order to demonstrate the scalability of the optimization and model comparison framework, we used the space of 12 models described below. We individually fitted the models and then performed model comparison to select the best fitting model. A set of null models were included which are anatomically implausible. If model selection performed correctly, then it is expected that these models would perform poorly.

To investigate the importance of known anatomical pathways in reconstructing the observed steady state statistics of the empirical local field potentials (i.e. autospectra and NPD), we considered a set of competing models. Specifically, we look at the role of four pathways and their interactions: the cortico-striatal indirect; the cortico-subthalamic hyperdirect; thalamocortical relay; and the subthalamic-pallidal feedback. In total we tested 6 families of models (presented later in the results section-figure 5):

1. + indirect.
2. + indirect/ + hyperdirect pathway.
3. + hyperdirect / - indirect.
4. + indirect / - hyperdirect/ + thalamocortical.
5. + indirect / + hyperdirect/ + thalamocortical.
6. -indirect / + hyperdirect/ + thalamocortical.

We considered these six families and divide them into two sub-families that do or do not include the subthalamopallidal (STN → GPe) feedback connection. Family (1) investigates whether the indirect pathway alone can explain the pattern of observed spectra and functional connectivity. In the case of family (2), previous work has highlighted the importance of hyper-direct connections in the functional connectivity (Jahfari et al., 2011; Nambu et al., 2015), yet anatomical research has shown dopamine to weaken the density of synaptic projections (Chu et al., 2017). Thus, family (2) provides an ideal set of models to examine the nonlinear mapping of anatomical to functional connectivity described in the introduction of this paper. Families (3) and (6) represent *null* models in which the indirect pathway is excluded and are used as implausible models to test the model comparison procedure. This is because it is thought that the indirect pathway is vital to explain activity within the network following dopamine depletion associated with PD (Albin et al., 1989; Alexander et al., 1986; Bolam et al., 2000). The functional role of the thalamocortical subnetwork is relatively unknown (but see recent work: Reis et al. 2019) and so families (4) and (5) provide an examination of whether the addition of the thalamic relay can better explain the empirical data. The second level of families (i.e. x.1-2) investigates whether the reciprocal network formed by the STN and GPe is required to explain observed patterns of connectivity in the data. This network has been the subject of much study and is strongly hypothesized to play an important role in the generation and/or amplification of pathological beta rhythms (Bevan et al., 2002; Cruz et al., 2011; Plenz and Kital, 1999).

## 3 Results

### 3.1 Properties of Iterative Fitting and Convergence when Applied to a Simple Model of the Pallido-Subthalamic Subcircuit

We provide an example model inversion in figure 2 to examine how the ABC algorithm iteratively converges to yield an approximation to the summary statistics of the empirical data. This example uses a simple model of a basal-ganglia subcircuit comprising the reciprocally connected subthalamic nucleus (STN) and external segment of the globus-pallidus (GPe) shown in figure 2A. Specifically, this model was fit to the autospectra and directed functional connectivity from the results originally reported in West et al., (2018) which described an analysis of local field potentials recorded from a rodent model of Parkinsonism (see methods for experimental details). By tracking the value of the objective function (i.e. the R^2^) over iterations (figure 2B) we demonstrate a fast-rising initial trajectory in the first 15 iterations that eventually plateaus towards convergence, that is well approximated by a logistic function (shown by purple dashed line). In figure 2C and D the simulated features (autospectra and NPD respectively) gradually move closer to the empirically estimated features with each iteration of the algorithm. Overall, average fits were R^2^ = 0.41 for the autospectra (figure 2C;) and R^2^ = 0.25 for the functional connectivity estimates (figure 2D).

**Figure 2.**
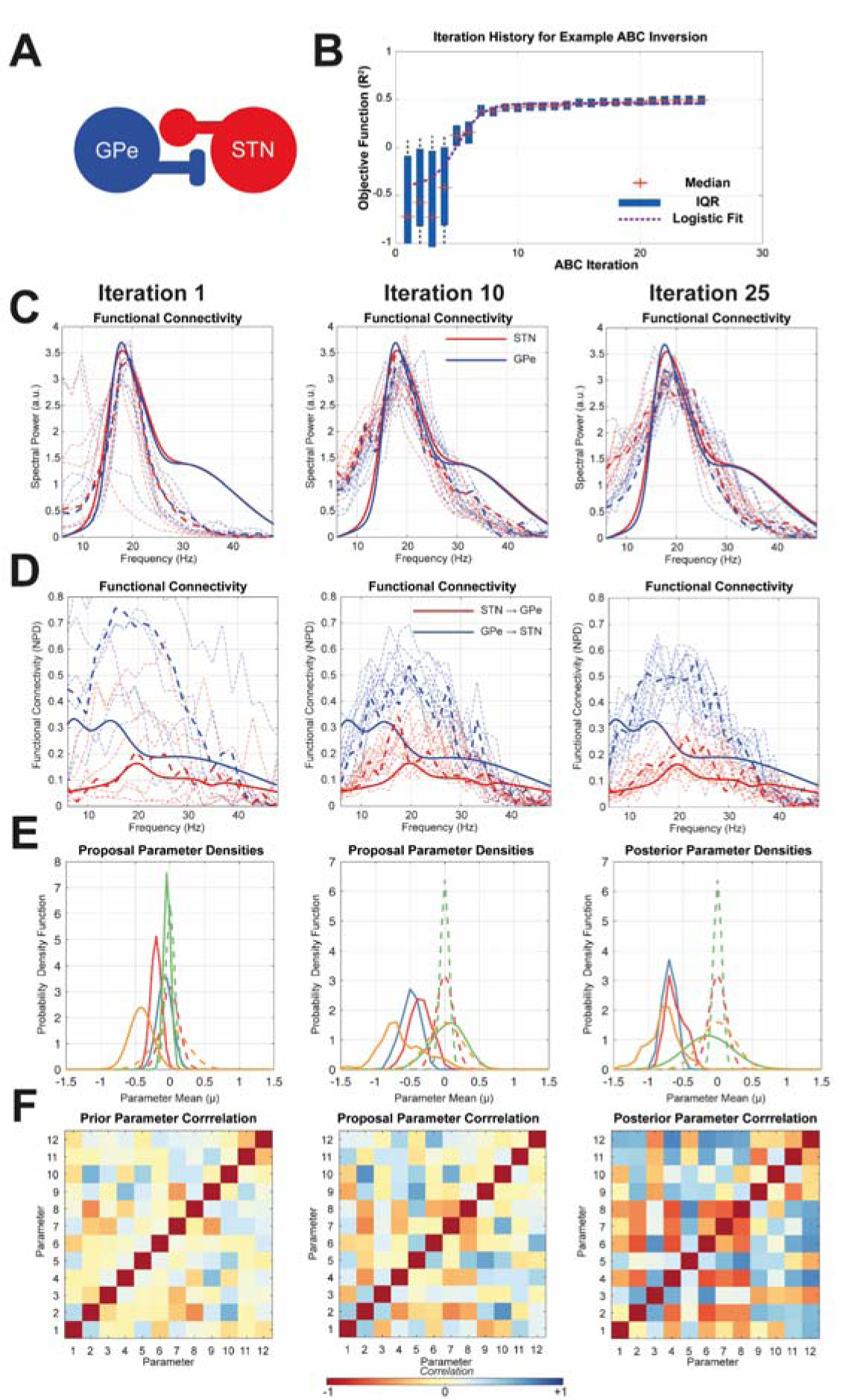
Examining the convergence of ABC optimization upon summary statistics from recordings of the STN and GPe in Parkinsonian rats. Parameters of neuronal state space models were optimized using the ABC method detailed in the text. Snapshots of the optimization are taken at the 1^st^, 10^th^, and 25^th^ iteration at which the optimization converges. **(A)** Schematic of the STN/GPe neural mass model. **(B)** The iteration history of the ABC algorithm is presented as a sequence of box plots indicating the distribution of fits (R^2^) at each sampling step, with mean and interquartile range indicated by individual crosses and boxes. Optimization shows a logistic convergence with a fit indicated by purple dashed line. **(C)** Autospectra of the empirical data (bold) and simulated data (dashed) are shown. The best fitting parameter sample for each iteration is given by the bold dashed line. **(D)** Similarly, the functional connectivity (non-parametric directionality; NPD) is shown in red and blue with the same line coding. **(E)** Examples of the prior (dashed) and proposal (bold) marginal distributions for a selection of five parameters are shown (note some priors have identical specifications and so overlap). It is seen that over iterations the proposal and posterior deviate from the prior as the latent parameter densities are estimated. **(F)** Correlation matrices from copula estimation of joint densities over parameters. Colour bar at bottom indicates the correlation coefficient. Correlated modes appear between parameters as optimization progresses.

The evolution of the proposed marginal densities (figure 2E) demonstrates that over the optimization, parameter means, and variances deviate significantly from the prior. Estimation of some parameters is better informed by the data than for others, as indicated by the different precision of the proposal densities (for example the parameter density outlined in blue exhibits reduced posterior precision). Additionally, learnt multivariate structure in the joint parameter densities is apparent from the correlation matrices of the t-Copula used in their estimation (see methods; figure 2F). The evolution of these matrices shows the emergence of distinct correlated modes.

The appearance of correlated modes acts to effectively reduce the dimensionality of the optimization problem: when estimating effective dimensionality of the parameter space by estimating the number of significant principal components (see methods) we find that optimized models show a reduction of 50-70% from that of the prior.

### 3.2 Testing Predictive Validity and Data Dependent Estimation of ABC Optimized Posteriors Using a Multi-start Procedure

This section of the results tests the predictive validity of the proposed framework i.e. that the scheme will make a consistent estimation of posterior model parameters across multiple realizations of optimization. This is achieved using the multi-start approach (Baritompa and Hendrix, 2005) described in the methods (section 2.7.1) in which we perform 10 instances of the algorithm on upon two separate datasets generated by different underlying models. The results of the multi-starts are shown in Figure 3.

**Figure 3.**
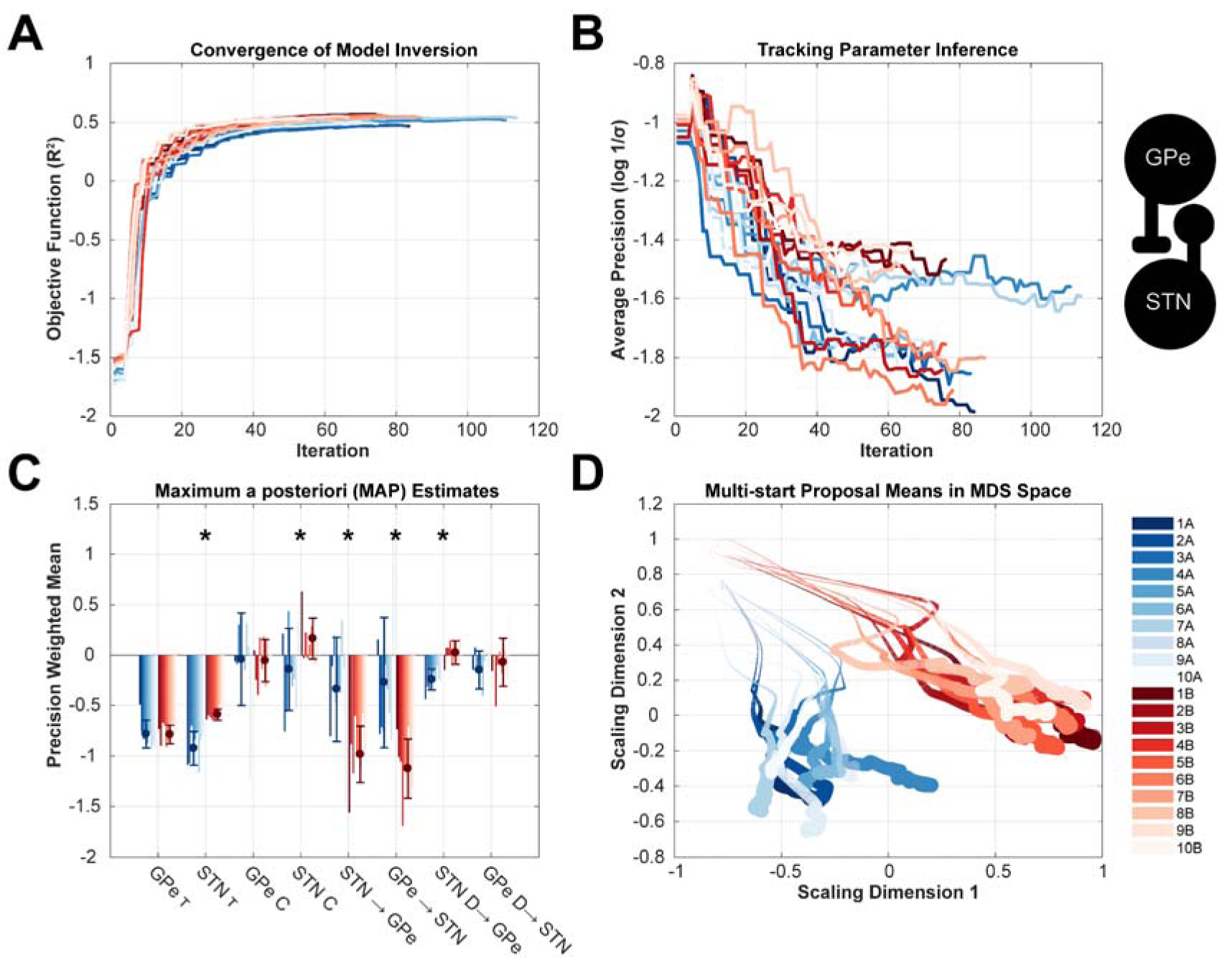
Multi-start analysis to test predictive validity of the ABC based estimation of model parameters by demonstrating consistency of estimation and the data specificity of parameter estimates. A two-node model of the STN/GPe circuit (inset) was fit to two different data sets: dataset *A* (blue) and dataset *B* (red) that were generated by different underlying models. Each estimation was performed 10 times with identical specification of prior distributions for all initializations. **(A)** Tracking of the objective function (R^2^) over the iterations demonstrated consistent convergence to 0.45 to 0.52 R^2^. **(B)** Optimization shows a consistent increase in the average precision (equivalent to a decrease in the logarithm of the inverse standard deviation of the data) of the posteriors indicating that data was informative in constraining parameter estimates. **(C)** Examination of the MAP estimates (weighted by their inverse precision) demonstrates a consistent inference of parameter values. Some parameters are drawn to common values with both data *A* and *B* (e.g. GPe time constant), whilst others show differences informed by the data (e.g. STN time constant). MAP values are given as log scaling parameters of the prior mean. The prior values were set to equal zero. Error bars give the standard deviations of the weighted estimates across initializations. Asterisks indicate significant t-test for difference in means between parameters estimated from data *A* and *B*. **(D)** To visualize trajectories of the multi-starts, the high dimensional parameter space was reduced to two dimensions using multi-dimensional scaling (MDS). Evolutions of the means of the proposal parameters exhibit a clear divergence between data sets A and B that were significantly different (MANOVA, see main text).

The evolution of the objective function (R^2^) over the progress of the ABC optimization is presented in figure 3A. The value of R^2^ reached by convergence is 0.55±0.02 for dataset A and 0.51±0.02 for dataset B showing consistency in the accuracy at convergence across the starts. In figure 3B, the average log-precision of the marginal densities is tracked over the progress of the optimization. These data show that across all initializations, the average precision of the posterior densities (1/σ = 0.025) was four times greater than those of the priors (1/σ = 0.1) demonstrating increased confidence in parameters estimates that were constrained by the data.

In figure 3C, we present the maximum a posterior (MAP) estimates for each parameter across the multi-starts. The posterior means were weighted by their precision (as described in the methods) such that parameters that were poorly informed by the data tended towards zero. In figure 3C, there are clear differences between parameters inverted upon the two separate sets of data (red versus blue bars; asterisks indicate significant t-tests). For instance, the mean STN time constant (2^nd^ group of bars from the left) is smaller for data A compared to B. Other parameters were well informed by the data, but not different between either data sets (e.g. GPe time constant; 1^st^ set of bars from the left). A third category of parameters were poorly inferred from the data (e.g. input gain to GPe; 3^rd^ set of bars from the left) where both estimates are close to zero deviation from the prior mean.

Finally, we apply statistical testing to the MAP estimates to determine (i) estimator consistency and (ii) differentiability with respect to different empirical data. To determine (i): estimator consistency, we applied a one-sample Hotelling test within a 10-fold, leave-one-out cross-validation to each set of MAP estimates from the multi-start. For both samples of parameters estimated from data A and B we find there to be a 0% rejection of the null hypothesis that the mean of the fold is significantly different from that of the left-out sample. This suggests that samples are consistent between initializations. To test (ii): that the posteriors were differentiated by the data on which they were estimated, we apply the Szekely and Rizzo energy test. We find there to be a significant difference in the means of the two samples (Φ = 6.68; P = 0.001). This finding is supported by a MANOVA test that demonstrates that the two data sets are significantly segregated by their posterior parameter means (D = 1, P < 0.001).

Visualization of the parameter space using multidimensional scaling (MDS) confirms the segregation of the posterior samples into two clusters determined by the datasets from which they are estimated. These results confirm that the ABC optimized posteriors are consistent across multiple initializations and that its output is determined by differences in the underlying model generating the given data.

### 3.3 Testing Face Validity of the Model Comparison Approach

To verify that the model comparison approach provides a valid identification of the original generative model of the data (c.f. face validity) we constructed a confusion matrix (as detailed in the methods section 4.6.2) using variations on the STN/GPe model presented in the previous sections and shown in figure 4A. In the case of validity, the analysis should correctly identify the system that originally generated the data (i.e. the best model scores should lay along the diagonal of the confusion matrix).

**Figure 4.**
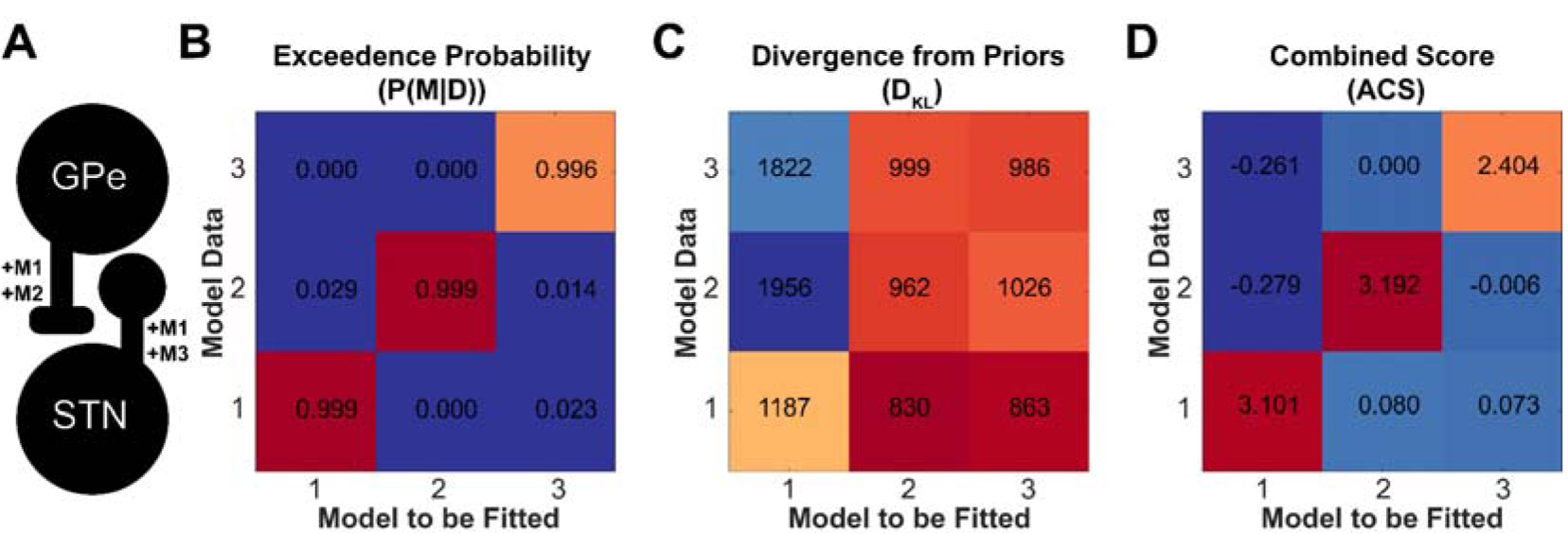
Testing face validity of the ABC model comparison approach to model identification. Confusion matrices were constructed by fitting the three models of the STN/GPe circuit. Synthetic data was generated using the fitted models and then the three original models were fitted back to the synthetic data to test whether model comparison could identify the generating model. **(A)** Schematic of neural mass model to be fitted. Annotations of connections indicate the presence of each for models 1-3. **(B)** Matrix of posterior model probabilities *P(M*|*D)* computed as exceedance probabilities across the joint model space. **(C)** Matrix of D_KL_ divergences of posteriors from priors. When *P*(*θ*|D_0_) = *P(*θ*)* then the divergence is zero thus larger values indicate a bigger shift from the prior. **(D)** Combined scoring to simultaneously account for model accuracies and divergence (ACS). Large values indicate better fits with parsimonious posteriors (small D_KL_).

In figure 4B we present the posterior model probabilities *P(M*|*D)* estimated by the exceedance probabilities computed from thresholding on model accuracy using a ϵ* computed from the joint model space (see methods). This analysis demonstrates that, in terms of accuracy, the most probable models lie on the diagonal of the confusion matrix showing that the comparison of models by their posterior accuracies is enough to correctly identify the generating model. In figure 4C, analyses show that the divergence of each model’s posteriors from priors (so called *complexity* measured in terms of the D_KL_) remain minimized along the diagonal such that best fitting models in panel A are not resulting from overfitting. In the case of model 1 (which is the most flexible in terms of numbers of free parameters) there are inflated divergences in the first column. These are the result of a large deviation of posteriors when attempting to fit the data generated from the alternative models. This shows that a post-hoc analysis of model parameter deviation (using D_KL_) can be used to discriminate models which have been overfitted. When combining these two measures into the ACS metric (summarising model accuracy minus complexity) in figure 4D, it is seen that the best fitting models are still correctly identified even when accounting for the increased complexity of posterior parameter densities. These results demonstrate that the model comparison approach can properly identify models from which the data originated, thus providing a face validation of the model comparison procedures.

### 3.4 Scaling up to Larger Model Spaces: Application to Models of the Cortico-Basal Ganglia-Thalamic Circuit

Finally, we apply the ABC framework to a larger and more complex model space to test the scalability of the methodology. Specifically, we devise a set of 12 models (illustrated in figure 5) incorporating combinations of pathways in the cortico-basal ganglia-thalamic circuit amongst a set of six neural populations motor cortex (M2); striatum (STR), GPe, STN, and thalamus (Thal.). Models are split into sets including/excluding the indirect (M2 → STR → GPe → STN); hyperdirect (M2 → STN); and thalamocortical relay (M2↔Thal.). Models are further subdivided to include or exclude the subthalmo-pallidal feedback connection (STN → GPe; models prefixed M *x*.2 to denote inclusion of the connection). For a full description and defence of model choices please see methods. These models were fit individually to the empirical data and then model comparison used to determine the best candidate model.

In figure 6 we show the resulting model fits and then the subsequent model comparison done in order to determine the best model or set of models. From visual inspection of the fits to the data features in figure 6A as well as the distribution of posterior model accuracies in figure 6B there is a wide range of model performances with regards to accurate fitting of the models to the data. Inspection of the posterior model fits to the data features in 6A shows that even in the case of the best fitting model (M 5.2) fitting fails to account for the multiple peaks in the autospectra, with models showing a preference for fitting the largest peak around 20 Hz. There is also a systematic overestimation of the directed functional connectivity (NPD) from cortex to subcortex. When the R^2^ of the fitted models are segregated between their autospectra and functional connectivity, the latter is more accurately fitted (M 5.2; R^2^ = 0.45) than for the power (M 5.2; R^2^ = −0.3).

**Figure 5.**
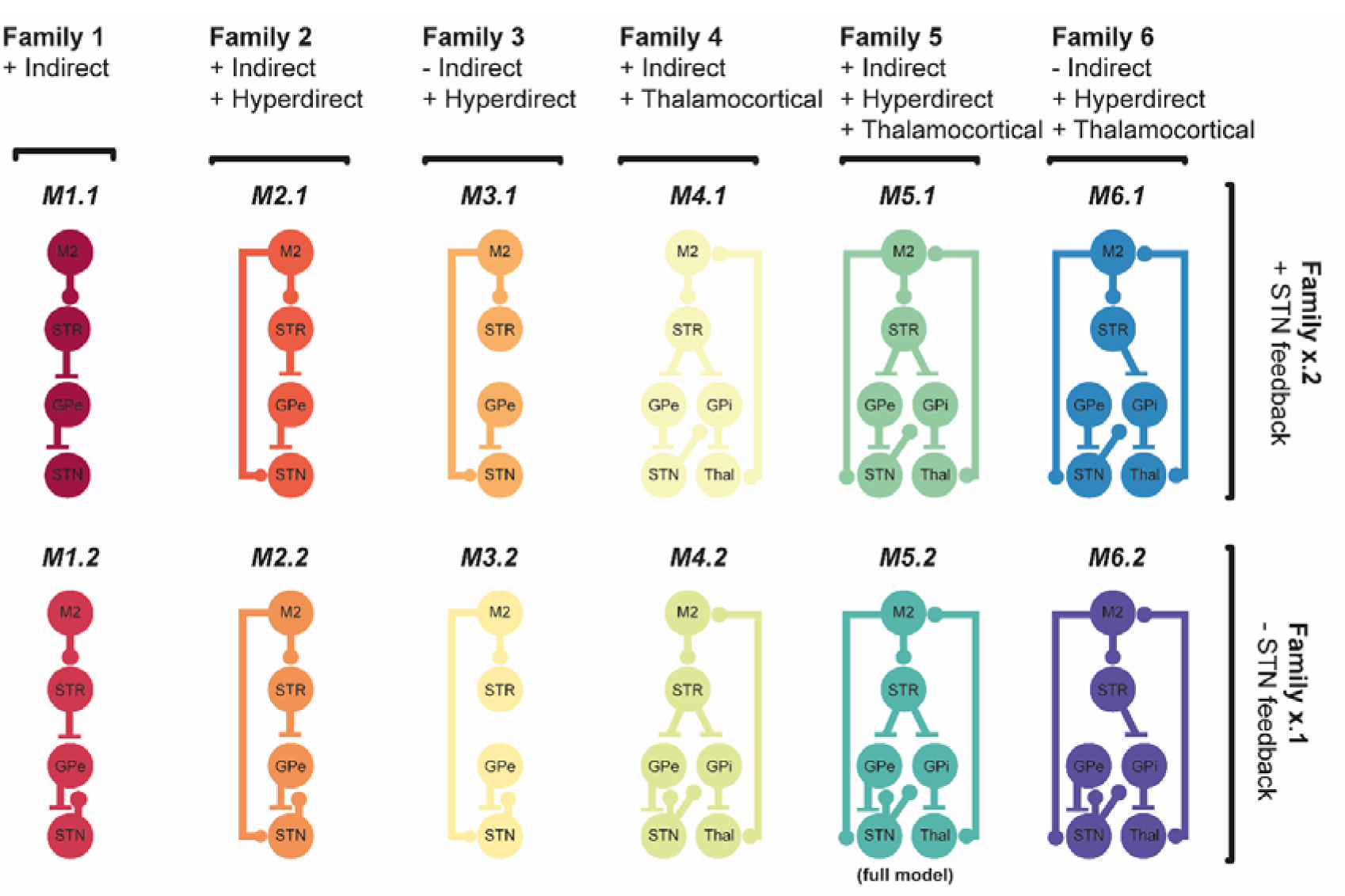
Illustration of the model space of the cortico-basal-ganglia network fitted with ABC and compared with Bayesian model selection. The model space comprises six families which can be further subdivided into two subfamilies yielding 12 models in total. Family (1) models the indirect pathway; family (2) contains models with both the indirect and hyperdirect pathways; family (3) contains models with the hyperdirect pathway but not indirect pathway; family (4) contains models with the indirect and thalamocortical relay; family (5) contains models with indirect, hyperdirect, and thalamocortical pathways; family (6) contains models with hyperdirect and thalamocortical pathway but indirect pathway. Finally, each family comprises two sub-families that either exclude (Mx.1) or include (Mx.2) subthalamopallidal feedback excitation. Excitatory projections are indicated by ball-ended connections, whilst inhibitory connections are flat-ended.

**Figure 6.**
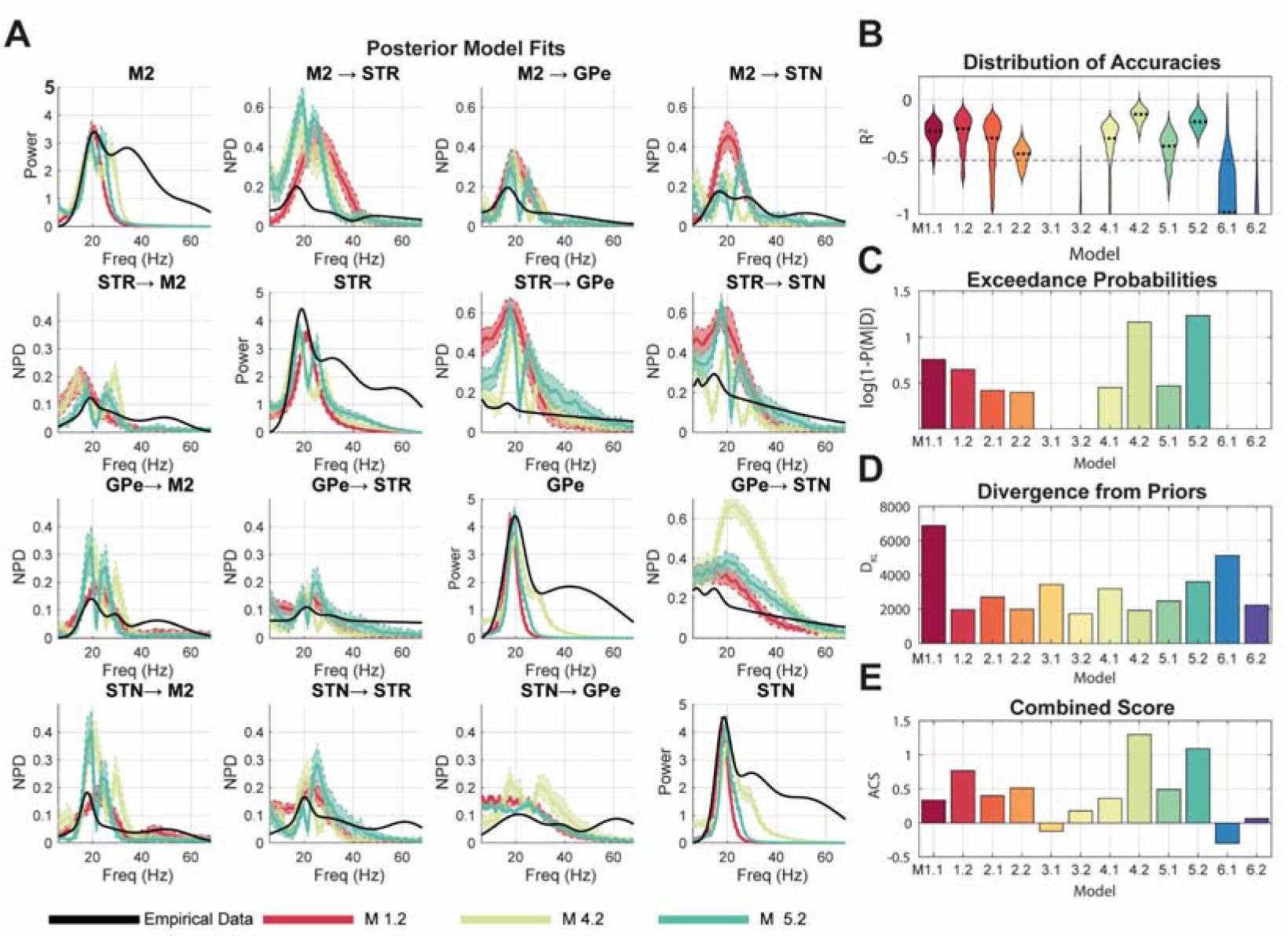
Scaling up the ABC model comparison framework – investigating models of the cortico-basal ganglia-thalamic network. 12 competing models (six families subdivided each into two sub-families) were fitted to empirical data from Parkinsonian rats. Models were fitted to summary statistics of recordings from the motor cortex (M2), striatum (STR), subthalamic nucleus (STN), and external segment of the globus pallidus (GPe). Models were first fit using ABC to estimate the approximate posterior distributions over parameters. To assess relative model performances, 1000 draws were made from each model posterior and corresponding data was simulated. **(A)** The posterior model fits for the top three performing models are shown, with autospectra on the diagonal and NPD on the off-diagonal (M 5.2 in light green; M 4.2 in turquois; and 1.2 in red). Bounds indicate the interquartile range of the simulated features. **(B)** Violin plots of the distributions of model accuracies (R^2^) of the simulated pseudo-data from the empirical data. **(C)** The exceedance probability approximation to the model evidence *P(M*|*D)* is determined by computing the number of samples from the posterior that exceed the median model accuracy (R^2^). **(D)** The Kullback-Leibler divergence of the posterior from prior is shown for each model. Large values indicate high divergence and overfitting. **(E)** Combined scores for accuracy and divergence from priors using ACS.

In all cases the models containing the subthalamopallidal excitatory connection (M *x*.2) perform better than those without (M *x*.1) and in good agreement with known Parkinsonian electrophysiology (Cruz et al., 2011). Notably we find that the model families 3 and 6 that both lack indirect pathway connections perform poorly with many of the posterior distributions of model fits falling far below the median of the whole model space. In the case of M 6.1 where there is a non-negligible exceedance probability, we see this accompanied by a high KL divergence and so both are penalized when measured with the combined ACS score. M 4.2 and 5.2 are the strongest models, with distributions of fits tightly clustered around high values yielding high model evidences. This suggests the importance of including thalamocortical feedback connections in the model. Scoring with ACS suggests model 4.2 is the best of the models due to the smaller model complexity (parameter divergence from prior). These results demonstrate the potential for the framework to be scaled up to investigate larger models and model spaces to investigate a neurobiological circuit. These posterior models can then be taken forward in post-hoc numerical simulations and/or analyses in order to explore their dynamics in further detail.

## 4 Discussion

### 4.1 Performance and Validity of ABC for Inverse Modelling of Brain Network Dynamics

In this paper we have formulated a framework for the inverse modelling of neural dynamics based upon an algorithm using ABC-SMC (Toni et al., 2009); (figure 1). We characterised the properties of this method when applied to models and data types typically used in systems neuroscience. Examination of the outcomes of the sequential ABC algorithm indicated that parameter estimates converge to yield best fit approximations to the summary statistics of empirical data and provide constraints upon the parameter estimates associated with a data dependent reduction in the estimated posterior precision (figure 2).

To assess the validity of the procedure we tested for both the face and predictive validity. Face validity was tested by building a confusion matrix from a set of synthetic data (figure 4). These results demonstrated that the model inversion and comparison approach are able to reliably identify the model that generated the data. These tests are similar to those performed for the validation of steady-state DCM (Moran et al., 2009). We also establish predictive validity (figure 3) using a multi-start procedure to demonstrate that parameter estimates are consistent across multiple inversions of the same data. Further we also established that model parameters are uniquely determined by the data features from which they have been estimated in a way that is consistent across multiple instances of the same model inversion. Finally, we demonstrated the scalability of the schema by applying it to a large set of candidate models describing the cortico-basal-ganglia-thalamic circuit (figure 5) and show feasibility of applying this method to biologically relevant problems (figure 6).

### 4.2 Comparison with Existing Techniques

The enthusiasm for inverse modelling approaches in systems neuroscience has led to an expansive effort to develop generic frameworks such as DCM that can be successfully applied across a range of modalities of electrophysiological data and models: spanning invasive animal recordings (e.g. Moran et al., 2011); electroencephalography (e.g. Boly et al., 2012); magnetoencephalography (e.g. Pinotsis et al., 2013); and recently calcium imaging (Rosch et al., 2018). In order to make these approaches widely applicable by making them computationally efficient, models are fit directly to the frequency-domain statistics of a neural time series (auto and cross spectral densities) by estimating the frequency response of the system around a stable equilibrium (Friston et al., 2014; Moran et al., 2009; Rowe et al., 2004). Fitting of models directly from their approximate frequency response at equilibrium enables a large reduction in the computational complexity of the problem by removing the need for numerical integration of the underlying equations of motion.

In this paper, we have introduced a framework which allows for the formal parameter estimation and model comparison of stochastic, non-equilibrium models of neural activity such that have typically been restricted to forward modelling approaches (e.g. Ritter et al., 2013). Recent work developing optimization of large scale neural models has shown that by improving parameter estimation algorithms, it is possible to fit high-dimensional, complex models of neural dynamics without appealing to steady-state approximations (Hadida et al., 2018). By relaxing the restrictions on model dynamics, it is thus possible to fit a model using a combination of a much wider range of data features beyond that contained in just the cross spectral density. Future work will be needed to identify potential candidate features to help better constrain parameter estimates.

By nature of the expansion of model behaviours that can be explored, the ABC framework comes with an increased computational demand over that of estimation from DCM. In the future, work should be done to characterize the specific demands and scalability of the problems that can be investigated within this framework. To provide a sense of time complexity, the evaluation of the model fitting and comparison presented in section 2.5 of the results took approximately 4 hours per model and approximately 30 hours for the complete analysis. We note however, the parallel nature of sampling steps would make the algorithm amenable to deployment on cluster computing such as that done in Aponte et al., (2016).

In future work, a test of construct validity should be conducted to examine the consistency of model estimates made between the ABC based method and existing approaches. Current methods adopting algorithms such as variational Laplace (Friston et al., 2007); deterministic sampling approaches (Hadida et al., 2018); and Generative Adversarial Networks (Arakaki et al., 2019) all provide potential routes to estimation of network mechanisms of spontaneous dynamics yet each is likely suited to different pairings of data and models. A quantitative assessment of their applicability in different scenarios would provide concrete answers to the inevitable question of which scheme is best is for a particular type of problem.

### 4.3 Addressing Limitations of ABC for Model Optimization and Modifications of the Framework

In recent years, ABC has become a well-established tool for parameter estimation in systems biology (Excoffier, 2009; Liepe et al., 2014; Ratmann et al., 2009; Toni et al., 2009; Turner and Sederberg, 2012). Nonetheless, it is known that ABC will not perform well in all cases and faces issues common across all model optimization problems (Sunnåker et al., 2013). Specifically, it has been shown that the simplest form of ABC algorithm based upon rejection-sampling approach is inefficient in the case where the prior densities lie far from the true posterior (Lintusaari et al., 2016). Fortunately, for biological models there is often a good degree of a priori knowledge as to the potential range of values in which a parameter is likely to fall. For instance, in the case of neural transmission delays we know that neural activity takes anywhere from a minimum of 1 ms up to a maximum of ∼ 50ms to propagate which allows the precision of prior parameter densities to be well constrained. This is a benefit of using biophysically rooted models over more abstract, phenomenological models where the range of potential parameter values are unknown.

As well as issues regarding uninformative priors, all optimization algorithms are subject to the so called ‘curse of dimensionality’ by which the exponential increase in the volume of the search space with each new parameter effectively means that any sampling procedure will in practice be extremely sparse. Furthermore, nonlinear models often exhibit non-convex, non-smooth objective functions in which multiple local minima exist, making the identification of the global minima difficult. In addition to highlighting the value of proper prior constraints on model parameters, we make use of two other modifications of the ABC procedure (Li et al., 2017) which aim to avoid the issue of ill-informed priors as well the local minima problem. The first relates to the non-parametric estimation of proposal densities using kernel density approximation. This property aids evaluation of non-convex objective functions during intermediate steps of the optimization as a) long tailed and skewed distributions can emerge to aid sampling when the true posterior lies far to the tails of the prior; and b) it facilitates simultaneous sampling of multiple regions over parameter space and thus can reduce the risk of converging towards a local extrema upon initialization. The second addition to the method employs copula estimation of the joint parameter densities from the non-parametric marginals (which are themselves estimated using kernel density approach). This facilitates the binding of parameters into correlated ‘modes’ that can reduce the effective volume of the parameter space to be searched, allowing for more highly parameterized models than would otherwise be possible with conventional ABC approaches (Li et al., 2017).

The selection of summary statistics are well known to be a vital factor in determining the outcomes of ABC estimated posterior (Beaumont et al., 2002; Sunnåker et al., 2013). The current approach does not consider the precision of the data features used to inform the model inversion. Such schemes exist for DCM where data features may be weighted in terms of an estimated noise term, a similar extension is likely to be of use with ABC inverted models, especially in the case where multiple types of summary statistics are combined.

It is also important to note that in the case of ABC for model selection, the usage of a Bayes factor derived from an approximate posterior model probability can be problematic in the case of adoption of an insufficient summary statistic (Robert et al., 2011). Necessary and sufficient conditions on summary statistics to ensure consistency in Bayesian model selection have been previously described (Marin et al., 2014). Despite this, approaches such as the validation performed here for model identification arguably provide the most flexible methods to determine the usage of a given statistic for data reduction.

Model comparison methods such as the one presented here have been criticized for covering only a finite hypothesis space and so ‘winning models’ are on the surface only the most plausible of a small selection of candidates (Lohmann et al., 2012; Templeton, 2009). It has been stated previously that this hypothetically infinite model space can be effectively reduced by specifying prior model probabilities (Friston et al., 2013), a decision implicit in the selection of a finite space of candidate models to be tested. This comes from a desire to test only models that may be expected to be reasonable *a priori*, and to ask hypotheses that are experimentally useful. For instance, in the model space presented in figure 6, we include models lacking STN → GPe feedback for purposes of demonstrating that the model comparison framework can identify models with poor anatomical grounding. In a real case of model comparison, we advocate a utilitarian selection of candidate models (Box, 1976) where including hypotheses where there is no strong *a priori* reasoning is not considered useful.

### 4.4 Conclusions

Overall, we have synthesised a framework for parameter estimation and model comparison that draws upon a number of recent developments in likelihood free, semi-parametric, inference that make it attractive to the inverse modelling of large-scale neural activity. This framework provides a robust method by which large scale brain activity can be understood in terms of the underlying structure of the circuits that generate it. This scheme avoids making appeals to local-linear behaviour and thus opens the way to future studies exploring the mechanisms underlying itinerant or stochastic neural dynamics. We have demonstrated that this framework provides consistent estimation of parameters over multiple instances; can reliably identify the most plausible model that has generated an observed set of data; and the potential for this platform to be scaled to larger models and datasets. Whilst this work constitutes a first validation and description of the method, more work will be required to establish its validity in the context of more complex models as well as statistics of non-stationary properties of neural dynamics.

## 5 Acknowledgments and Funding

T.O.W. acknowledges funding from UCL CoMPLEX doctoral training program, and the UCL Bogue Fellowship. H.C. receives funding from an MRC Career Development award (MR/R020418/1). S.F.F. receives funding support from the UCL/UCLH NIHR Biomedical Research Centre. L.B. acknowledges funding support from the Leverhulme Trust (RPG-2017-370). The Wellcome Trust Centre for Neuroimaging is funded by core funding from the Wellcome Trust (539208). We thank Peter Magill and Andrew Sharott at the Brain Network Dynamics Unit, Oxford University for making available the experimental data used in this study. We thank all authors of the publicly available toolboxes used in this paper (listed in the supplementary information VI).

## Notes

### Competing Interest Statement

The authors have declared no competing interest.

### Summary of Updates

Major revision of text for clarity.

