## Supplementary Information for "Model Based Inference of Large Scale Brain Networks with Approximate Bayesian Computation"

### 1 Table of Methods Used

| Method | Notes | Reference(s) |
| --- | --- | --- |
| Convolution based neural mass models | Mean field approximation to homogenous neural population activity. | [1–3] |
| Neural mass model of the cortico-basal ganglia -thalamic circuit | Population model with structure and parameterization defended and explained in given references. | [4,5] |
| Likelihood free inference with Approximate Bayesian Computation | References are for introductions/tutorials | [6–8] |
| Sequential Approximate Bayesian Computation | Improvement on ABC to aid convergence and computational efficiency | [9,10] |
| Kernel density approximation to ABC marginals and copula estimation of dependence | We use a cross-validation log-likelihood optimization of the kernel density bandwidth for estimation of the marginals, with approximate maximum likelihood to fit a t-copula to estimate the joint. | [11] |
| Model Selection with ABC Optimized Models |  | [12,13] |
| Electrophysiological recordings from Parkinsonian rats | Field Recordings made in experimental 6-OHDA model of Parkinsonism | [4,14] |
| Non-parametric Directionality for directed functional connectivity estimates | Used as a summary statistic of between signal interactions. | [15,16] |

### 2 Simulation of Neural Model

##### 2.1.1 Notation

We denote spike rates by *x*, membrane potentials by *v*, synaptic gains by *H*, time constants by $\tau$, and external inputs by *A*. First order derivatives are indicated using Newton’s notation. Subscripts denote the *i*-th population except in the case of sigmoid operators *S* and their coefficients *R* for which a second within-population index *q* is given if there is more than one operator per population such as in models with multiple interacting populations (i.e. cortex) or populations with self-inhibition. Throughout we will use a plain letter for a vector (e.g. y) and a letter with a subscript for a vector element (e.g. y_i_).

##### 2.1.2 Modelling of Neural Dynamics

In order to simulate neural signals generated by neuronal populations, we used a coupled network of neural mass models that simulate field potentials generated synchronized activity of large ensembles of neurons [2,17]. A single neural mass approximates the population activity of a large number of homogenous neurons [18]. These models rest on the assumption that an incoming volley of spikes arriving at a population can be converted to a post synaptic potential by convolution with a synaptic response kernel. Following temporal integration of the post synaptic potentials, the population can then in turn generate a spike density with a sigmoidal mapping of membrane voltage to spike output frequency. Populations may be coupled via *intrinsic* connectivity to simulate the dynamics within cortical columns or *extrinsic* connectivity to simulate inter-areal coupling [3,19]. A single mass of coupled populations is given generally by the form:

$\dot{v_{i}}=x_{i}$,

$$\dot{\boldsymbol{x}_{\boldsymbol{i}}}\boldsymbol{=}\frac{\boldsymbol{H}_{\boldsymbol{i}}}{\boldsymbol{\tau}_{\boldsymbol{i}}}\left( \boldsymbol{u}_{\boldsymbol{i}}\boldsymbol{+}\boldsymbol{S}_{\boldsymbol{i}}\left( \boldsymbol{A}_{\boldsymbol{i}} \right) \right)\boldsymbol{-}\frac{\boldsymbol{2}}{\boldsymbol{\tau}_{\boldsymbol{i}}}\boldsymbol{x}_{\boldsymbol{i}}\boldsymbol{-}\frac{\boldsymbol{1}}{\boldsymbol{\tau}_{\boldsymbol{i}}^{\boldsymbol{2}}}\boldsymbol{v}_{\boldsymbol{i}}\boldsymbol{,}$$

Equation 1

where the average postsynaptic membrane potential of the $i$th population is given by $v_{i}$, and is parameterized by a synaptic gain $H_{i}$ and a lumped post synaptic time constant $\tau_{i}$. The input to the mass is given between the outer set of brackets and comprises some background noise $u_{i}$ plus a combined input from the other *J* populations. To couple distant populations, the contributions to population *i* from *j* are weighted using adjacency matrix $\omega,$ and summed across all *J* populations:

$$A_{i}=\sum_{j=1}^{J} \omega_{ij}v_{j}$$

Equation 2

Masses within separate cortical columns are intrinsically coupled such that inhibitory and excitatory populations interact. Inhibitory cells have negative connection weights and vice versa for excitatory cells.

The final terms of equation 1. relate to the fact that these equations are equivalent to a convolution operation of an exponential kernel (see Jansen and Rit 1995 for details of the derivation). Membrane potentials are converted to spike densities via the sigmoid operator:

$$S_{i}(A_{i})=1/(1+e^{-R_{i}A_{i}}),$$

Equation 3

which is parameterised by $R^{i}$ to determine the slope of the activation function (a parameter specific to population *i*) and effectively models the variance of the population’s firing thresholds.

To create the full model describing the basal ganglia and motor cortex, masses are coupled with a structure outlined in the schematic in the full model (M 5.2) shown in figure 6. We model inhibitory connections by flipping the sign on the adjacency matrix, such that they have a subtractive influence. Connectivity was simulated between sources (extrinsic connectivity), the neural activity (given by *V*) propagates between sources according to a weighted adjacency matrix with entries A indicating the presence and sign of connections. The adjacency matrix of the full model is given below

$$\omega=\left[ \begin{matrix} 0 & 0 & 0 & 0 & 0 & \omega_{1,6} \\ \omega_{2,1} & 0 & 0 & 0 & 0 & 0 \\ 0 & \omega_{3,2} & 0 & \omega_{3,4} & 0 & 0 \\ \omega_{4,1} & 0 & \omega_{4,3} & 0 & 0 & 0 \\ 0 & \omega_{5,2} & 0 & \omega_{5,4} & 0 & 0 \\ 0 & 0 & 0 & 0 & \omega_{6,5} & 0 \end{matrix} \right]$$

Equation 4

where column 1 gives connections projecting from M2; column 2 from the STR; column 3 from the GPe; column 4 from the STN; column 5 from the GPi; and column 6 from the thalamus. Equivalently the rows give the weights of the input to the populations. Variants of this full model can then be created by adjusting the parameters or removing coefficients $\omega_{ji}$ from the matrix.

#### 2.2 Transmission Delays

We incorporate finite transmission delays by formulating the state space equations to explicitly depend on the past values of the sending node. This is unlike the approximations used in previous studies using DCM [20]. This was achieved by modifying the extrinsic connectivity matrices by indexing past values with a matrix $\boldsymbol{D}$ with elements $D_{ij}$ specifying the delay for connection of population $j$ to $i$. Thus, the total external input to node $i$ at time $t$ is:

$$A_{i}\left( t \right)=\sum_{j=1}^{J} \omega_{ij}v_{j}\left( t-D_{ij} \right),$$

Equation 5

with the constraint that $D_{ij}>0$.

#### 2.3 Integration of Stochastic Differential Equations

The model also incorporates a stochastic input $u_{i}$for each population which represents endogenous background activity. This input is given by:

$$u_{i}=C_{i}W,$$

Equation 6

where

$$W\sim N\left( 0,\sigma\right),$$

Equation 7

and $C_{i}$ represents a gain factor on the noise scaling the noise for population $i$. The noise $W$is drawn from a zero-mean normal distribution, with a standard deviation $\sigma$ that is set for the whole model. In the model presented here the stochastic innovations are independent of the state variable. In this case, the interpretation of the stochastic integral is simplified, and forward Euler with a suitably small step size (less than half of the fastest time constant) has been demonstrated to yield accurate results [21]. For the purposes of integration, we rescaled the noise by the square-root of the integration step $h$ to ensure stochastic fluctuations are smaller than the deterministic parts of the equation (as per Hansen et al. 2006):

$\hat{u_{i}}=u_{i}\surd h$.

Equation 8

It is important to note that the naïve application of deterministic integration schemes to stochastic problems has been demonstrated to lead to inaccurate or at worst, spurious, solutions [21]. Furthermore, approximations to absorb delay equations (in order to simplify the integration of the equations) using numerical approximation of the system’s Jacobian [23,24], as is done in dynamic causal modelling is inaccurate when applied to the same types of neural mass models presented here [25]. To allow for settling of state equations, we set the initial states to be equal to zero, and then remove the initial transient.

### 3 State Equations of Full Model

In total there are 14 state equations that are an adaption of the full model described in van Wijk et al. (2018). Each mass comprises two equations (decomposing the second order differential equations into two first order ones) that describes the voltage change of the population.

The motor cortex model consists of 4 populations of neurons and 8 state equations. Each layer is connected via ‘intrinsic’ (within source) connectivity with synaptic gain parameters (*H*). All subpopulations in the motor cortex have a self-inhibiting connection. In order to notate the *z^th^* intrinsic connections of the *i^th^* population, we extend our subscripts for synaptic gains to *H_i,z_*. All layers receive independent stochastic inputs *u_i_*. The cortical source comprises:

1) A middle layer composed of middle pyramidal cells with inhibitory self-connection (with strength parameterized by H_1_):

$\dot{v_{1}}=x_{1}$,

$\dot{x_{1}}=\frac{1}{\tau_{1}}\left( -H_{1,1} S_{1}\left( v_{1} \right)-H_{1,3} S_{1}\left( v_{3} \right)+H_{1,8} S_{1}\left( v_{2} \right)+S_{1}(A_{1})+u_{1} \right)-\frac{2}{\tau_{1}}x_{1}-\frac{1}{\tau_{1}^{2}}v_{1}$;

Equation 9

2) A supra-granular layer composed of superficial pyramidal cells with inhibitory self-connection (with strength parameterized by H_7_):

$\dot{v_{2}}=x_{2}$,

$\dot{x_{2}}=\frac{1}{\tau_{2}}\left( {-H}_{1,7} S_{2}\left( v_{2} \right)+H_{1,2} S_{2}\left( v_{1} \right)-H_{1,12} S_{2}\left( v_{3} \right)+H_{1,14} S_{2}\left( v_{4} \right)+u_{2} \right)-\frac{2}{\tau_{2}}x_{2}-\frac{1}{\tau_{2}^{2}}v_{2}$;

Equation 10

3) The supra-granular layer also contains a separate inhibitory interneuron population, again with an inhibitory self-connection (with strength parameterized by H_4_):

$\dot{v_{3}}=x_{3}$,

$\dot{x_{3}}=\frac{1}{\tau_{3}}\left( {-H}_{1,4} S_{3}\left( v_{3} \right)+H_{1,5} S_{3}\left( v_{1} \right)+H_{1,6} S_{3}\left( v_{4} \right)+H_{1,13} S_{3}\left( v_{2} \right)+u_{3} \right)-\frac{2}{\tau_{3}}x_{3}-\frac{1}{\tau_{3}^{2}}v_{3}$;

Equation 11

4) Finally, the infra-granular layer is made up of deep pyramidal cells also with an inhibitory self-connection (with strength parameterized by H_10_)

$\dot{v_{4}}=x_{4}$,

$\dot{x_{4}}=\frac{1}{\tau_{4}}\left( {-H}_{1,10} S_{4}\left( v_{4} \right)-H_{1,9} S_{4}\left( v_{3} \right)+H_{1,11} S_{4}\left( v_{2} \right)+u_{4} \right)-\frac{2}{\tau_{4}}x_{4}-\frac{1}{\tau_{4}^{2}}v_{4}$.

Equation 12

Overall the output of the cortex is equal to the voltage in the deep pyramidal layer, thus:

$V_{1}=v_{4}$.

Equation 13

The striatal source (STR) is modelled as a single inhibitory population and self-inhibitory connection (with strength parameterized by H_11_):

$\dot{v_{5}}=x_{5}$,

$\dot{x_{5}}=\frac{1}{\tau_{5}}\left( {-H}_{5,1} S_{5}\left( v_{5} \right)+S_{5}(A_{5})+u_{5} \right)-\frac{2}{\tau_{5}}x_{5}-\frac{1}{\tau_{5}^{2}}v_{5}$;

Equation 14

and total output:

$V_{2}=v_{5}$.

Equation 15

The source modelling the external segment of the pallidus (GPe) source is taken to be a single inhibitory population and self-inhibitory connection (with strength parameterized by H_12_):

$\dot{v_{6}}=x_{6}$,

$\dot{x_{6}}=\frac{1}{\tau_{6}}\left( S_{6}\left( A_{6} \right)+u_{6} \right)-\frac{2}{\tau_{6}}x_{6}-\frac{1}{\tau_{6}^{2}}v_{6}$,

Equation 16

and total output:

$V_{3}=v_{6}$.

Equation 17

The subthalamic nucleus (STN) source is modelled as a single excitatory population:

$\dot{v_{7}}=x_{7}$,

$\dot{x_{7}}=\frac{1}{\tau_{7}}\left( S_{7}(A_{7})+u_{7} \right)-\frac{2}{\tau_{7}}x_{7}-\frac{1}{\tau_{7}^{2}}v_{7}$,

Equation 18

and total output:

$V_{4}=v_{7}$.

Equation 19

The internal segment of the pallidus (GPi) source is taken to be a single inhibitory population and self-inhibitory connection (with strength parameterized by H_13_):

$\dot{v_{8}}=x_{8}$,

$\dot{x_{8}}=\frac{1}{\tau_{8}}\left( S_{8}\left( A_{8} \right)+u_{8} \right)-\frac{2}{\tau_{8}}x_{8}-\frac{1}{\tau_{8}^{2}}v_{8}$,

Equation 20

and total output:

$V_{4}=v_{8}$.

Equation 21

The thalamic population (Thal.) source is taken to be a single excitatory population with self-inhibition (with strength parameterized by H_14_):

$\dot{v_{9}}=x_{9}$,

$\dot{x_{9}}=\frac{1}{\tau_{9}}\left( -H_{9,1}S_{9}\left( v_{9} \right)+S_{9}\left( A_{9} \right)+u_{9} \right)-\frac{2}{\tau_{9}}x_{9}-\frac{1}{\tau_{9}^{2}}v_{9}$,

Equation 22

and total output:

$V_{5}=v_{9}$.

Equation 23

### 4 Table of Prior Parameter Values

| Parameter | Mean Value(s) | Units | Log Precision |
| --- | --- | --- | --- |
| Cortical Model |  |  |  |
| Intrinsic Connectivity:  $\boldsymbol{H}_{\boldsymbol{1,i}}$ | [400 800 400 400 400 400 400 400 400 400 800 400 400 400] | s^-1^ | 1/4 σ^2^ |
| Time Constants:  $\boldsymbol{\tau}_{\boldsymbol{1\ldots4}}$ | [3 2 12 18] | ms | 1/4 σ^2^ |
| Sigmoid Slope: $\boldsymbol{S}_{\boldsymbol{1\ldots4}}$ | All 2/3 |  | 1/4 σ^2^ |
| STR Model |  |  |  |
| Self-Inhibition:$\boldsymbol{H}_{\boldsymbol{5}}$ | 400 | s^-1^ | 1/8 σ^2^ |
| Time Constant: $\boldsymbol{\tau}_{\boldsymbol{5}}$ | 8 | ms | 1/8 σ^2^ |
| Sigmoid Slope:$\boldsymbol{S}_{\boldsymbol{5}}$ | All 2/3 |  | 1/4 σ^2^ |
| GPe Model |  |  |  |
| Self-Inhibition: | 0 | s^-1^ | 1/8 σ^2^ |
| Time Constant: $\boldsymbol{\tau}_{\boldsymbol{6}}$ | 8 | ms | 1/8 σ^2^ |
| Sigmoid Slope:$\boldsymbol{S}_{\boldsymbol{6}}$ | 2/3 |  | 1/8 σ^2^ |
| STN Model |  |  |  |
| Self- Inhibition: | 0 | s^-1^ | 1/8 σ^2^ |
| Time Constant: $\boldsymbol{\tau}_{\boldsymbol{7}}$ | 4 | ms | 1/8 σ^2^ |
| Sigmoid Slope:$\boldsymbol{S}_{\boldsymbol{7}}$ | 2/3 |  | 1/8 σ^2^ |
| GPi Model |  |  |  |
| Self- Inhibition: | 0 | s^-1^ | 1/8 σ^2^ |
| Time Constant: $\boldsymbol{\tau}_{\boldsymbol{8}}$ | 8 | ms | 1/8 σ^2^ |
| Sigmoid Slope:$\boldsymbol{S}_{\boldsymbol{8}}$ | 2/3 |  | 1/8 σ^2^ |
| Thal. Model |  |  |  |
| Self-Inhibition:  $\boldsymbol{H}_{\boldsymbol{9}}$ | 400 | s^-1^ | 1/8 σ^2^ |
| Time Constant: $\boldsymbol{\tau}_{\boldsymbol{9}}$ | 8 | ms | 1/8 σ^2^ |
| Sigmoid Slope:$\boldsymbol{S}_{\boldsymbol{9}}$ | 2/3 |  | 1/8 σ^2^ |
| Delays |  |  |  |
| M2 → STR: $\boldsymbol{D}_{\boldsymbol{5,4}}$ | 3 | ms | 1/4 σ^2^ |
| M2 → STN: $\boldsymbol{D}_{\boldsymbol{7,4}}$ | 3 | ms | 1/4 σ^2^ |
| STR → GPe: $\boldsymbol{D}_{\boldsymbol{6,5}}$ | 7 | ms | 1/4 σ^2^ |
| STR → GPi: $\boldsymbol{D}_{\boldsymbol{8,5}}$ | 12 | ms | 1/4 σ^2^ |
| GPe → STN: $\boldsymbol{D}_{\boldsymbol{7,6}}$ | 1 | ms | 1/4 σ^2^ |
| GPe → GPi: $\boldsymbol{D}_{\boldsymbol{8,6}}$ | 1 | ms | 1/4 σ^2^ |
| STN → GPe:$\boldsymbol{D}_{\boldsymbol{6,7}}$ | 3 | ms | 1/4 σ^2^ |
| STN → GPi: $\boldsymbol{D}_{\boldsymbol{8,7}}$ | 3 | ms | 1/4 σ^2^ |
| GPi → Thal.: $\boldsymbol{D}_{\boldsymbol{9,8}}$ | 3 | ms | 1/4 σ^2^ |
| Thal. → M2: $\boldsymbol{D}_{\boldsymbol{1,9}}$ | 3 | ms | 1/4 σ^2^ |
| M2 → Thal.: $\boldsymbol{D}_{\boldsymbol{9,4}}$ | 8 | ms | 1/4 σ^2^ |
| Connections |  |  |  |
| M2 → STR: $\boldsymbol{A}_{\boldsymbol{5,4}}$ | (+) 2000 | s^-1^ | 1/4 σ^2^ |
| M2 → STN: $\boldsymbol{A}_{\boldsymbol{7,4}}$ | (+) 2000 | s^-1^ | 1/4 σ^2^ |
| STR → GPe: $\boldsymbol{A}_{\boldsymbol{6,5}}$ | (-) 1600 | s^-1^ | 1/4 σ^2^ |
| STR → GPi: $\boldsymbol{A}_{\boldsymbol{8,5}}$ | (-) 1600 | s^-1^ | 1/4 σ^2^ |
| GPe → STN: $\boldsymbol{A}_{\boldsymbol{7,6}}$ | (-) 2000 | s^-1^ | 1/4 σ^2^ |
| GPe → GPi: $\boldsymbol{A}_{\boldsymbol{8,6}}$ | (-) 2000 | s^-1^ | 1/4 σ^2^ |
| STN → GPe:$\boldsymbol{A}_{\boldsymbol{6,7}}$ | (+) 2000 | s^-1^ | 1/4 σ^2^ |
| STN → GPi: $\boldsymbol{A}_{\boldsymbol{8,7}}$ | (+) 2000 | s^-1^ | 1/4 σ^2^ |
| GPi → Thal.: $\boldsymbol{A}_{\boldsymbol{9,8}}$ | (-) 1600 | s^-1^ | 1/4 σ^2^ |
| Thal. → M2: $\boldsymbol{A}_{\boldsymbol{1,9}}$ | (+) 1000 | s^-1^ | 1/4 σ^2^ |
| M2 → Thal. : $\boldsymbol{A}_{\boldsymbol{9,4}}$ | (+) 2000 | s^-1^ | 1/4 σ^2^ |

### 5 Computation of Kullback-Leibler Divergence for Multivariate Normal Distribution

In order to compute the full multivariate divergence between posterior $P_{1}=P\left( \theta| D_{0} \right)=N(\mu_{1},\Sigma_{1})$ and prior distributions $P_{0}=P\left( \theta| D_{0} \right)= N(\mu_{0},\Sigma_{0})$ over parameters we make an approximation to a *k* dimensional multivariate normal distribution and compute the Kullback-Leibler distance evaluated from the mean and covariance of the distributions:

$$KL\left( P_{1} | |P_{0} \right)=\frac{1}{2}\left( tr(\Sigma_{0}^{-1}\Sigma_{1})+\left( \mu_{0}-\mu_{1} \right)^{T}\Sigma_{1}^{-1}\left( \mu_{0}-\mu_{1} \right)-k+ln\left( \frac{det\Sigma_{0}}{det\Sigma_{1}} \right) \right)$$

Equation 24

### 6 List of Toolboxes Used

We thank all authors of the toolboxes below:

| Toolbox Name | Author | Year | Source/Reference |
| --- | --- | --- | --- |
| allcomb | ‘Jos’ | 2018 | <https://uk.mathworks.com/matlabcentral/fileexchange/10064-allcomb-varargin> |
| boundedline-pkg | Kelly Kearney | 2015 | <https://github.com/kakearney/boundedline-pkg> |
| bplot | Jonathan C. Lansey | 2015 | <https://uk.mathworks.com/matlabcentral/fileexchange/42470-box-and-whiskers-plot-without-statistics-toolbox> |
| brewermap | Stephen Cobeldick | 2014 | <https://github.com/DrosteEffect/BrewerMap> |
| export_fig | Oliver J. Woodford, Yair M. Altman | 2014 | <https://github.com/altmany/export_fig> |
| highdim | Brian Lau | 2017 | <https://github.com/brian-lau/highdim> |
| hotellingT2 | Antonio Trujillo-Ortiz | 2002 | <https://uk.mathworks.com/matlabcentral/fileexchange/2844-hotellingt2> |
| linspecer | Jonathan C. Lansey | 2015 | <https://github.com/davidkun/linspecer> |
| neurospec 2.2 | David Halliday | 2018 | <http://www.neurospec.org/> |
| ParforProgMon | Dylan Muir, Willem-Jan de Goeij, The MathWorks, Inc. | 2016 | <https://github.com/DylanMuir/ParforProgMon> |
| splitvec | Bruno Luong | 2009 | <https://uk.mathworks.com/matlabcentral/fileexchange/24255-splitvec> |
| violin | Holger Hoffmann | 2015 | <https://uk.mathworks.com/matlabcentral/fileexchange/45134-violin-plot> |

### 7 Auxiliary References

8.	Sisson SA, Fan Y, Beaumont M. ABC Samplers. Handbook of approximate Bayesian computation. Boca Raton: Chapman and Hall/CRC; 2018.
